# The first gapless, reference-quality, fully annotated genome from a Southern Han Chinese individual

**DOI:** 10.1101/2022.08.08.503226

**Authors:** Kuan-Hao Chao, Aleksey V Zimin, Mihaela Pertea, Steven L Salzberg

**Affiliations:** Department of Computer Science, Johns Hopkins University, Baltimore, MD 21218, USA; Center for Computational Biology, Johns Hopkins University, Baltimore, MD 21218, USA; Department of Biomedical Engineering, Johns Hopkins University, Baltimore, MD 21218, USA; Department of Biostatistics, Johns Hopkins University, Baltimore, MD 21211, USA

**Keywords:** genome assembly, annotation, DNA sequencing, reference genome, variant calling

## Abstract

We used long-read DNA sequencing to assemble the genome of a Southern Han Chinese male. We organized the sequence into chromosomes and filled in gaps using the recently completed CHM13 genome as a guide, yielding a gap-free genome, Han1, containing 3,099,707,698 bases. Using the CHM13 annotation as a reference, we mapped all genes onto the Han1 genome and identified additional gene copies, generating a total of 60,708 genes, of which 20,003 are protein coding. A comprehensive comparison between the genes revealed that 235 protein-coding genes were substantially different between the individuals, with frameshifts or truncations affecting the protein-coding sequence. Most of these were heterozygous variants in which one gene copy was unaffected. This represents the first gene-level comparison between two finished, annotated individual human genomes.

## Introduction

Over the past twenty years, the biomedical research community has relied on a single human reference genome, first published in 2001 (International Human Genome Sequencing Consortium 2001; Venter *et al*. 2001) when it was less than 90% complete and steadily improved since then. That reference genome, currently version GRCh38, is a mosaic of many individuals, which means that it does not capture the actual genome of any single human. In addition, the GRCh38 genome has hundreds of gaps, including very large gaps for all of the centromeres and the short arms of acrocentric chromosomes. In 2022, a breakthrough publication from the Telomere-to-Telomere (T2T) consortium described the first-ever complete human genome, CHM13, in which all gaps had been filled and which represented a single individual of northern European descent (Nurk *et al*. 2022). Although CHM13 describes a female and is missing the Y chromosome, the T2T consortium finished a gap-free Y chromosome from another individual, HG002, and released that as part of the CHM13 assembly.

The existence of a finished human genome, together with highly accurate long-read sequencing technology, now allows the assembly of additional human genomes of similarly high quality and completeness. As demonstrated first by the assembly of an Ashkenazy individual, Ash1 (Shumate *et al*. 2020), and more recently by the assembly of a Puerto Rican individual, PR1 (Zimin *et al*. 2021), long-read data (either on its own or in combination with short reads) can be used to generate highly accurate contigs covering most of the human genome, and these can then be scaffolded using either GRCh38 or CHM13 as a guide. Following scaffolding, the CHM13 sequence can then be used to fill in all gaps, including the centromeres, to produce a gap-free assembly.

An individual genome is far more useful as a research resource if it is annotated. Several computational pipelines can produce annotation *de novo* for a newly assembled genome, but when the genome represents another individual human, as opposed to a new species, *de novo* annotation is unnecessary. Instead, one can simply map (or “lift over”) the annotation from a reference human genome onto the new assembly. This is the approach we followed.

The first genome sequenced from a Han Chinese individual (designated YH), which was also the first Asian individual genome assembled, was published over a decade ago (Wang *et al*. 2008). Subsequently, another southern Han Chinese individual genome, HX1, was *de novo* assembled using single-molecule real-time (SMRT) long reads (Shi *et al*. 2016). Research then shifted toward northern individuals: the first northern Han Chinese individual genome, NH1.0, appeared in 2019 (Du *et al*. 2019), and the most recent Han Chinese haplotype-resolved genome, HJ-H1 and HJ-H2, was released in early 2022 (Yang *et al*. 2022).

In addition to these four Han Chinese genomes, two other individual genomes, each representing an ethnic group closely related to Han, have been assembled: ZF1 (a Tibetan individual) (Yang *et al*. 2022) and TJ1.p0/TJ1.p1 (a Tujia individual) (Lou *et al*. 2022). However, all of these genomes still contain gaps and none were fully annotated. Table 1 provides a comparison among these genomes and the new Han1 assembly.

**Table 1.**
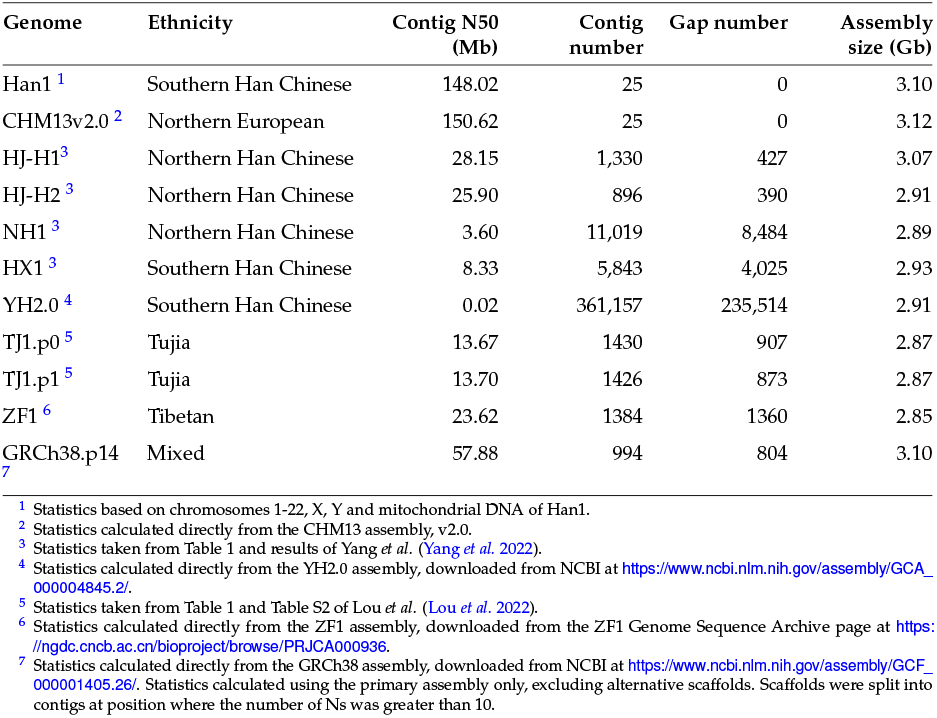
A comparison among Han1, GRCh38, and the assemblies of previously released Chinese genomes. Assembly sizes include estimated gaps.

| Genome | Ethnicity | Contig N50 (Mb) | Contig number | Gap number | Assembly size (Gb) |
| --- | --- | --- | --- | --- | --- |
| Han1 <sup>1</sup> | Southern Han Chinese | 148.02 | 25 | 0 | 3.10 |
| CHM13v2.0 <sup>2</sup> | Northern European | 150.62 | 25 | 0 | 3.12 |
| HJ-H1 <sup>3</sup> | Northern Han Chinese | 28.15 | 1,330 | 427 | 3.07 |
| HJ-H2 <sup>3</sup> | Northern Han Chinese | 25.90 | 896 | 390 | 2.91 |
| NH1 <sup>3</sup> | Northern Han Chinese | 3.60 | 11,019 | 8,484 | 2.89 |
| HX1 <sup>3</sup> | Southern Han Chinese | 8.33 | 5,843 | 4,025 | 2.93 |
| YH2.0 <sup>4</sup> | Southern Han Chinese | 0.02 | 361,157 | 235,514 | 2.91 |
| TJ1.p0 <sup>5</sup> | Tujia | 13.67 | 1430 | 907 | 2.87 |
| TJ1.p1 <sup>5</sup> | Tujia | 13.70 | 1426 | 873 | 2.87 |
| ZF1 <sup>6</sup> | Tibetan | 23.62 | 1384 | 1360 | 2.85 |
| GRCh38.p14 <sup>7</sup> | Mixed | 57.88 | 994 | 804 | 3.10 |

<sup>1</sup> Statistics based on chromosomes 1-22, X, Y and mitochondrial DNA of Han1.
<sup>2</sup> Statistics calculated directly from the CHM13 assembly, v2.0.
<sup>3</sup> Statistics taken from Table 1 and results of Yang *et al.* (Yang *et al.* 2022).
<sup>4</sup> Statistics calculated directly from the YH2.0 assembly, downloaded from NCBI at https://www.ncbi.nlm.nih.gov/assembly/GCA_000004845.2/.
<sup>5</sup> Statistics taken from Table 1 and Table S2 of Lou *et al.* (Lou *et al.* 2022).
<sup>6</sup> Statistics calculated directly from the ZF1 assembly, downloaded from the ZF1 Genome Sequence Archive page at https://ngdc.cncb.ac.cn/bioproject/browse/PRJCA000936.
<sup>7</sup> Statistics calculated directly from the GRCh38 assembly, downloaded from NCBI at https://www.ncbi.nlm.nih.gov/assembly/GCF_000001405.26/. Statistics calculated using the primary assembly only, excluding alternative scaffolds. Scaffolds were split into contigs at position where the number of Ns was greater than 10.

Here we describe the assembly of deep-coverage long-read data to produce the genome of a Southern Han Chinese male individual, which we designate Han1. We filled in all gaps using CHM13 sequence where necessary, and then annotated the genome by mapping over the latest RefSeq annotation from CHM13 onto Han1. We then compare the annotation between Han1 and CHM13 and report on some of the key differences.

## Materials and methods

### Sample background

The data for the HG00621 genome were generated and made publicly available by the Human Pangenome Reference Consortium (https://github.com/human-pangenomics/hpgp-data). HG00621 is a Southern Han Chinese male from Fu Jian Province located on the southeastern coast of China. We chose HG00621 for this study because it represents a population for which no annotated genome previously existed, and because Han is the largest ethnic group in the world. The estimated population size of Han people is approximately 1.4 billion, with around 1.29 billion living in China (National Bureau of Statistics of China 2021) and 21.8 million living in Taiwan (Central Intelligence Agency 2022). Moreover, as the largest population located in East Asia, the Han group plays a key role in understanding ancient human migrations, since it is on the path connecting Africa to the Pacific Islands (Cavalli-Sforza 1998; Zhang *et al*. 2007).

Han Chinese are divided into two population groups, the northern Han (NH) and southern Han (SH), a division strongly supported by genetic markers as well as dental, craniometric and archaeological evidence (Tongmao *et al*. 1987; Zhao and Dao Lee 1989; Chu *et al*. 1998; Cavalli-Sforza 1998). One of the reasons that we chose a southern Han Chinese is because most Han immigrants to the United States, Taiwan, Singapore, Malaysia, Philippines, and other territories originated from southern China (Cavalli-Sforza 1998). A complete and accurate Han Chinese reference genome should be a valuable resource for genetic studies in this population. Another benefit of selecting HG00621 is that sequencing data are available from both of his parents, HG00619 and HG00620, which provides opportunities to study allelic variations in the future. Also worth noting is that at least three of HG00621’s grandparents are confirmed Southern Han Chinese individuals.

### Genome assembly

Our main goal for this study was to create a gapless, reference-quality, haplotype-merged assembly of the HG00621 genome with 25 sequences representing chromosomes 1 to 22, X, Y, and the mitochondrial DNA. We assembled contig sequences *de novo*, and we used the CHM13 genome recently published by the T2T consortium to guide the scaffolding process. The CHM13 genome is far more complete and accurate than the previous standard, GRCh38 (Nurk *et al*. 2022). CHM13 represents a northern European individual, and because it is a female sample it lacks a Y chromosome. We used version 1.1 of the CHM13 assembly (https://github.com/marbl/CHM13), a gap-free version augmented with the complete Y chromosome from HG002, an individual of Ashkenazi Jewish descent.

The initial Han1 assembly was created using PacBio high-fidelity (HiFi) reads and Oxford Nanopore Technology (ONT) ultra-long reads, with approximately 39x coverage in HiFi reads and 35x coverage in ONT reads (Table 2). We assembled HiFi reads with Hifiasm (Cheng *et al*. 2021) version 0.16.1-r375 and ONT reads with Flye (Kolmogorov *et al*. 2019) version 2.5. The statistics of these two preliminary assemblies are shown in Table 3. Comparing both assemblies, the HiFi draft assembly had substantially higher contiguity with an N50 contig size of 95.8 Mb and only 182 contigs, while the Flye assembly had an N50 of 40.9 Mb and 1,658 contigs. We therefore chose the Hifiasm output as the main assembly for the subsequent steps.

**Table 2.** Sequencing data from HG00621 used for assembly of the Han1 genome.

| Sequence type | Number of reads | N50 read length (bp) | Mean read length (bp) | Maximum length | Total length (Gbp) | Genome coverage |
| --- | --- | --- | --- | --- | --- | --- |
| PacBio HIFI | 5,570,675 | 21,499 | 21,989 | 50,388 | 122.50 | 39.45x |
| ONT Ultralong | 3,110,293 | 83,822 | 34,136 | 2,495,296 | 106.17 | 34.75x |

**Table 3.** Statistics for the preliminary Han1 assemblies.

| Assembler | Sequencing data | Assembled sequence | Contig N50 | Number of contigs |
| --- | --- | --- | --- | --- |
| Hifiasm | PacBio HiFi | 3,110,501,483 | 95,769,069 | 182 |
| Flye | ONT "ultralong" | 2,974,205,132 | 40,850,737 | 1,658 |

For the scaffolding step, we first used the MaSuRCA chromosome scaffolder module (Zimin *et al*. 2017) (‘chromosome_scaffolder.sh’) with the CHM13 reference to order and orient contigs generated by Hifiasm. This process is similar to the scaffolding procedures we used in building the Ashkenazi (Ash1) (Shumate *et al*. 2020) and Puerto Rican (PR1) (Zimin *et al*. 2021) reference genomes. The chromosome scaffolder identified 10 misassembled contigs that we had to split before scaffolding.

After scaffolding, we had 25 scaffolds (with gaps) representing nuclear chromosomes 1-22, X and Y, plus 77 unplaced contigs containing an additional 38,815,834 bp. The chromosome sequences contained only 101 gaps.

We then proceeded with three gap closing steps. For these steps we used MaSuRCA’s intra-scaffold gap-closer script (‘close_scaffold_gaps.sh’), which is a wrapper for the SAMBA scaffolder (Zimin and Salzberg 2022). In the first step we used Hi-Fi reads and default parameters, which closed 24 gaps. The second step used contigs generated by Flye from the ONT reads with modified parameters ‘-m 2500 -o 10000’, which set the thresholds for the minimum match length on both sides of a gap (2,500bp) and the maximum overhang (10,000bp). This step closed an additional 19 gaps. The third and final step of automated gap closing used the CHM13 sequence to attempt closing the remaining 58 gaps. This step closed another 45 gaps with only 8 gaps remaining in chromosomes 13, 14, 15 and 22. Manual inspection of these gaps identified repeat-induced misassemblies in pericentromeric regions in chromosomes 13, 14 and 15, which we corrected manually, and redundant haplotype-variant contigs in chromosomes 15 and 22 that we deleted. After fixing the misassemblies and eliminating redundant haplotype contigs, we re-ran gap closing and ended up with a single contig for every chromosome. We observed that one of the unplaced contigs contained several copies of the mitochondrial sequence, and we extracted a single circular copy using alignment to the mitochondrial sequence from GRCh38.

We screened all 77 unplaced contigs for contamination by using KrakenUniq (Breitwieser *et al*. 2018) to align them to a database containing bacteria, viruses, and common vectors. We identifed one contig as the Epstein-Barr virus, a common DNA viral contaminant in human DNA. The size of this contig was 177,625 bp, which is close to the 170 Kb genome size of Epstein–Barr virus (Chang *et al*. 2009). This contig was removed. We aligned all remaining unplaced contigs to the nuclear chromosomes and found that all were redundant copies of variant haplotypes.

The last step of the process was to polish the chromosome sequences with the HiFi reads using JASPER (Guo *et al*. 2022) version 1.0.0 to correct errors and reduce reference bias in the patches of sequence that originated from the Flye assembly and CHM13 genome. This step also produced a quality value (QV) estimate of 57 for the polished assembly, which corresponds to fewer than 2 errors per megabase.

To compare the Han1 and CHM13 genomes, we aligned Han1 to CHM13 using Nucmer (Marçais *et al*. 2018), selected the best 1-to-1 alignments using delta-filter (Kurtz *et al*. 2004), and created dotplots using mummerplot.

### Gene annotation

We used Liftoff (Shumate and Salzberg 2021) version 1.6.2 to map genes from CHM13 onto Han1. The CHM13 annotation was generated by mapping RefSeq (O’Leary *et al*. 2016) release 110 of GRCh38.p14 onto CHM13 version 2.0. The CHM13 annotation contains a total of 61,140 genes, including 20,022 protein-coding genes and 18,389 lncRNA genes. It includes 181,713 transcripts of which 129,878 are protein-coding. The CHM13 annotation used here is available at ftp://ftp.ccb.jhu.edu/pub/data/T2T-CHM13/.

Although Liftoff maps most genes automatically, the ribosomal DNA (rDNA) regions present a problem because they occur in very long tandem arrays of near-identical copies. Each rDNA unit is composed of three ribosomal RNA (rRNA) genes, 18S, 5.8S, and 28S, separated by two flanking spacers (5’- and 3’-ETS) and two internal transcribed spacers (ITS-1 and 2) (Agrawal and Ganley 2018). The arrays are located on the acrocentric chromosomes 13, 14, 15, 21 and 22. To accurately map the rDNA arrays onto Han1, we adopted a 2-pass Liftoff process.

First we ran Liftoff to map all genes except the rDNA arrays onto Han1, as follows. We aligned a 44,838bp reference rDNA sequence from the KY962518 locus (Kim *et al*. 2018) to Han1 using Nucmer (Marçais *et al*. 2018) with the parameters: ‘ –maxmatch -l 31 -c 100’, which identified 1196 preliminary rDNA locations. We then filtered these matches to retain only those that were >10,00bp and >98.5% identical to the reference rDNA sequence, which yielded 257 approximate rDNA array locations. We used bedtools (Quinlan and Hall 2010) maskfasta to mask the locations of the 257 potential rDNA arrays, replacing them with Ns, and then ran Liftoff with parameters: ‘-chroms <chroms_mapping.csv> -copies -polish’ to map all CHM13 genes except the rDNA arrays. This Liftoff first-pass procedure mapped 60,099 genes, including 20,003 protein-coding genes, 18,321 lncRNAs, and 51 non-rDNA array rRNAs, as well as many other gene types including microRNAs and pseudogenes.

In our second pass process, we only lifted over the rDNA annotations from CHM13 onto Han1. We first extracted all rDNA annotations on CHM13 into an annotation (GFF) file, which included 219 rDNA units with 3 subunits each, and thus 657 subunits in total. Then we ran Liftoff with parameters: ‘-f <features.txt> -mm2_options “-N 250” -copies -sc 0.95’. The <features.txt> restricted the mapping to locations where all three subunits mapped onto Han1 at the same locus in the correct order, 18S, 5.8S, and 28S, which we called a valid rDNA unit lift-over condition. One additional note is that the rDNA morphotypes on each chromosome are structurally distinct (Nurk *et al*. 2022) which can create a mapping bias. We checked all 219 rDNA arrays during the mapping process to ensure that each copy was mapped to the best-matching chromosome. The Liftoff second-pass procedure mapped a total of 203 rDNA units, which is 609 subunits in total.

To verify the rDNA mapping result, we intersected the mapped annotations from the first and second passes using bedtools (Quinlan and Hall 2010) intersect version 2.30.0 with parameters: ‘-wa -wb’. None of the 203 rDNA units overlap with any other genes mapped to Han1, which shows no sign of mismapped rDNA units. In the final step, we merged the two GFF files and ran gffread (Pertea and Pertea 2020) version 0.12.7 with parameters: ‘-O -F –keep-exon-attrs’ to sort the final annotation and set the phases of CDS features.

### CHM13 and Han1 annotation comparison

We compared the gene annotations between CHM13 and Han1 by running LiftoffTools (Shumate and Salzberg 2022) version v0.2.0, which performed a transcript-level comparison between all transcripts on the two genomes. For protein-coding transcripts, LiftoffTools determines the effects of any DNA differences on the protein sequence, which it categorizes into 10 groups: synonymous, nonsynonymous, inframe deletion, inframe insertion, start codon loss, 5’ truncation, 3’ truncation, frameshift and stop codon gain.

Liftofftools reported comparisons between 181,029 transcripts between CHM13 and Han1, with only 720 transcripts unmapped. For 130,909 transcripts with coding sequence (CDS) features (which included protein-coding genes, VDJ segments, and some pseudogenes), this step computed DNA sequence identity, amino acid sequence identity, and any of the ten variant effects mentioned above. For the 50,120 transcripts lacking CDS features, only the DNA sequence identity was computed.

To analyze mutations at the gene level, we mapped 130,909 transcripts back to their gene names and obtained a list of 19,706 protein-coding genes, 25 C regions, 75 J segments, and 403 V segments, for a total of 20,209 genes. For each of these, we selected the transcript with the highest amino acid sequence identity score as representative, and categorized the gene into the variant effect group of that transcript. The number of genes in each group is summarized in Table 4.

**Table 4.** Comparison of genes with coding sequence (CDS) features mapped from CHM13 to Han1. For this gene-level analysis, the transcript-to-transcript mapping selected the alignment that produced the highest amino acid identity score for the coding sequence. Truncated genes are those where the mapped copy is missing either the 5’ end or the 3’ end of the transcript, including the start/stop codon.

| Mapped genes with CDS features |  |  |  |  |  |  |  |  |
| --- | --- | --- | --- | --- | --- | --- | --- | --- |
| Conserved genes (82%) |  | Altered Genes (18%) |  |  |  |  |  |  |
| Identical | synonymous | non-synonymous | start lost | stop gained | truncated | frameshift | in-frame insertion | in-frame deletion |
| 14130 | 2442 | 3200 | 21 | 31 | 1+39<br>(5'+3') | 143 | 110 | 92 |

We were particularly interested in genes containing deleterious mutations. We assumed that non-synonymous mutations, in-frame insertions, and in-frame deletions are relatively harmless, and we focused on mutations that created major differences between the CHM13 and Han1 protein sequences, which included frameshifts (an insertion or deletion whose length is not a multiple of 3), truncations that removed either the 5’ or 3’ end of the transcript including part of the coding sequence, mutations that changed the start codon to another codon, and mutations that created a premature stop codon. This gave us a target list of 235 genes for further investigation.

Because many of the 235 genes with damaging mutations might represent cases where the Han1 genome is heterozygous, and where one haplotype contains a normal copy of the gene, we went back to the raw HiFi data and used it to identify heterozygous and homozygous variants, as follows. We first aligned all HiFi reads to the CHM13 genome using Minimap2 (Li 2018) version 2.24-r1122, and then used samtools (Li *et al*. 2009) version 1.13 to sort and index the aligned BAM file. We called variants using the mpileup and call commands from bcftools (Li 2011) version 1.15.1, and selected heterozygous and homozygous variants by running bcftools filter with parameters ‘GT=“het”‘ and ‘GT=“hom”‘ respectively. This step identified 4,331,520 heterozygous sites and 2,285,451 homozygous sites.

We manually inspected a sample of mutated genes with heterozygous mutations, and found in each case that the alternative haplotype lacked the damaging mutation. Therefore we focused the remainder of our analysis on genes with homozygous mutations. We filtered out single nucleotide polymorphisms (SNPs) and kept mutations that might contribute to frameshifts, leaving us with 204,706 homozygous mutation sites.

Next, we extracted all CDS regions from all transcripts of the 235 genes with the most damaging mutations, which yielded a total of 8,578 coding exons. We intersected these with the 204,706 homozygous, non-SNP mutation sites using bedtools intersect. After removing repeats caused by exon sharing between isoforms, 54 distinct mutation sites in 46 distinct genes were identified, where each gene contained at least 1 homozygous, non-SNP mutation as compared to CHM13.

### Protein-level analysis of mutated genes

We further investigated our list of 46 mutated genes. We first checked the reported homozygous mutation sites in IGV (Robinson *et al*. 2011), extracted the GFF entries with target gene names, and translated their coding sequences (CDS) into amino acid sequences using gffread (Pertea and Pertea 2020). We then compared the protein sequence of Han1 and CHM13 to RefSeq using Blastp (Altschul *et al*. 1997) and parasail (Daily 2016) to align the sequences and calculate the amino acid sequence identity.

### Gene copy number analysis

We compared copy number variation between Han1 and CHM13 by grouping genes into paralogous groups using Liftofftools, which uses the ‘linclust’ function in MMSeq2 (Steinegger and Söding 2017). Protein-coding genes were clustered based on their amino acid sequences, and non-coding genes were clustered based on their nucleotide sequences, where genes in a cluster needed to be at least 90% identical over 90% of their lengths to at least one other cluster member.

## Results and discussion

### Genome assembly

The Han1 genome (version 1.0) contains 3,099,707,698 bp, which includes the 22 autosomes, chromosomes X and Y, and the mitochondrial genome. Approximately 120Mb in gaps was filled using sequence from the CHM13 genome, illustrated in Figure 1. The sizes of each chromosome and the amount of non-HG00621 sequence per chromosome are summarized in Table 5. Han1 is slightly smaller than CHM13, which has a total size of 3.117 Gbp. Chromosomes 9 and Y have the lowest proportion of HG00621 sequence compared to other chromosomes. Chromosome 9 has the largest centromeric region among all autosomes, as has been previously reported (Humphray *et al*. 2004). The centromeric region spans approximately 35 Mb, which was inserted entirely from CHM13. Chromosome Y contains a number of particularly challenging regions for assembly, including the male-specific region, MSY, which is composed of eight very long palindromes ranging from 9 Kb (P7) to 1.45 Mb (P1) with arm-to-arm sequence identities of 99.94 - 99.997% (Skaletsky *et al*. 2003; Rozen *et al*. 2003), as well as a lengthy X-transposed region, XTR, sharing 99% sequence identity to chromosome X (Skaletsky *et al*. 2003; Ross *et al*. 2005), which led us to rely on the CHM13 assembly for 27% of chromosome Y. Note that the CHM13 Y chromosome was derived from a separate individual, HG002, of Ashkenazi Jewish descent.

**Figure 1.**
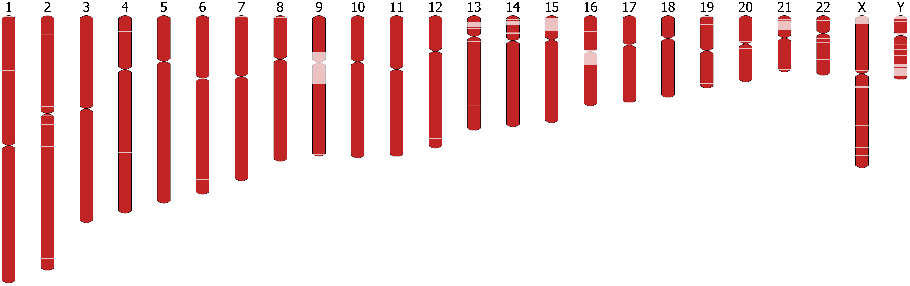
A visualization of the 24 chromosomes in Han1. Regions in red were assembled from HG00621, and regions in light pink are sequences inserted from CHM13. Graphics created using ideogram.js (https://github.com/eweitz/ideogram).

**Table 5.**
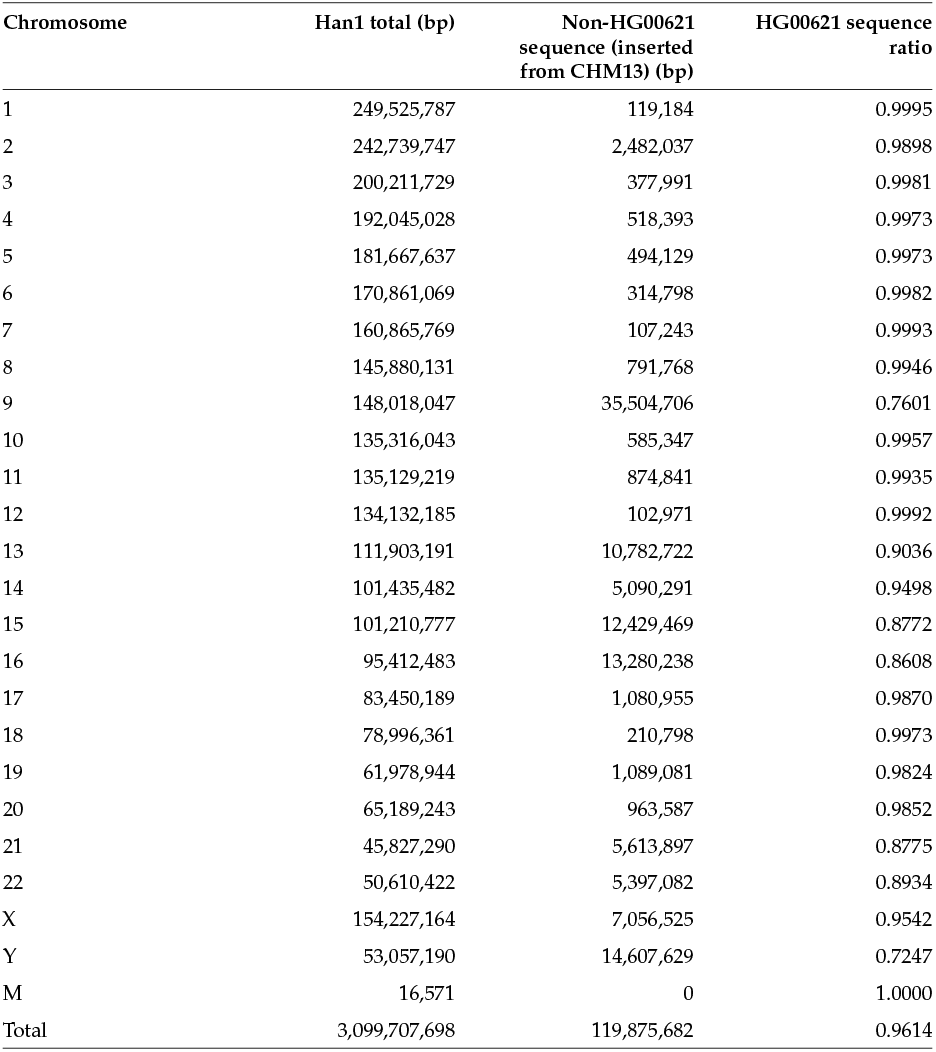
Han1 chromosome sizes and the amount of non-HG00621 sequence per chromosome in the final Han1 assembly. Every chromosome is in a single contig.

| Chromosome | Han1 total (bp) | Non-HG00621<br>sequence (inserted<br>from CHM13) (bp) | HG00621 sequence<br>ratio |
| --- | --- | --- | --- |
| 1 | 249,525,787 | 119,184 | 0.9995 |
| 2 | 242,739,747 | 2,482,037 | 0.9898 |
| 3 | 200,211,729 | 377,991 | 0.9981 |
| 4 | 192,045,028 | 518,393 | 0.9973 |
| 5 | 181,667,637 | 494,129 | 0.9973 |
| 6 | 170,861,069 | 314,798 | 0.9982 |
| 7 | 160,865,769 | 107,243 | 0.9993 |
| 8 | 145,880,131 | 791,768 | 0.9946 |
| 9 | 148,018,047 | 35,504,706 | 0.7601 |
| 10 | 135,316,043 | 585,347 | 0.9957 |
| 11 | 135,129,219 | 874,841 | 0.9935 |
| 12 | 134,132,185 | 102,971 | 0.9992 |
| 13 | 111,903,191 | 10,782,722 | 0.9036 |
| 14 | 101,435,482 | 5,090,291 | 0.9498 |
| 15 | 101,210,777 | 12,429,469 | 0.8772 |
| 16 | 95,412,483 | 13,280,238 | 0.8608 |
| 17 | 83,450,189 | 1,080,955 | 0.9870 |
| 18 | 78,996,361 | 210,798 | 0.9973 |
| 19 | 61,978,944 | 1,089,081 | 0.9824 |
| 20 | 65,189,243 | 963,587 | 0.9852 |
| 21 | 45,827,290 | 5,613,897 | 0.8775 |
| 22 | 50,610,422 | 5,397,082 | 0.8934 |
| X | 154,227,164 | 7,056,525 | 0.9542 |
| Y | 53,057,190 | 14,607,629 | 0.7247 |
| M | 16,571 | 0 | 1.0000 |
| Total | 3,099,707,698 | 119,875,682 | 0.9614 |

We identified two nuclear mitochondrial sequences (NUMTs) (Lopez *et al*. 1994) in Han1. The first one, an 866 bp insertion on chromosome 13 from 12,339,933 to 12,340,799 is present in CHM13, but absent from GRCh38. The second NUMT is a longer 13,781bp insertion on chromosome 20 from 21,541,750 to 21,555,531, and it is unique to the Han1 genome, i.e. it is absent from both CHM13 and GRCh38. We validated these two NUMTs by aligning the ultra-long ONT reads back to the Han1 genome using minimap2 with the parameter ‘–secondary=no’, requiring that the reads contained the entire NUMT and that the alignments extended at least 500bp beyond it on both ends. We found 15 ONT reads supporting the NUMT on chromosome 13, suggesting that it may be a heterozygous insertion, and 26 ONT reads supporting the chromosome 20 NUMT. The chromosome 20 NUMT includes 26 genes of which 8 are protein-coding genes, 16 are tRNAs, and 2 are rRNAs.

A dot plot comparing the entire sequence of Han1 and CHM13 genomes is shown in Figure 2, which illustrates the overall high collinearity of all chromosomes. One interesting finding is that Han1 contains an inversion at chr8: 7,306,407-11,410,283 (4,103,876 bp) which corresponds to CHM13 chr8: 11,717,452-7,617,994 (4,099,458 bp). This inversion is one of the largest common inversion polymorphisms in humans (Logsdon et al. 2021), and the arrangement in Han1, shown in more detail in Figure 3, matches GRCh38 but disagrees with CHM13. A more detailed dotplot of the inversion is shown in Supplementary Figure S1. This region includes the *β*-defensin gene cluster locus, and has been described as one of the most structurally dynamic regions in the human genome (Ganz 2003; Hollox et al. 2003, 2008; Mohajeri et al. 2016).

**Figure 2.**
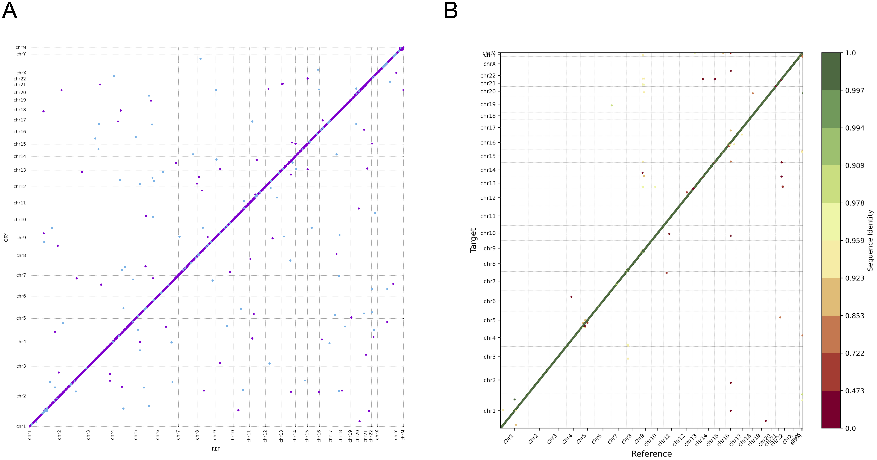
(A) A dot plot between the DNA sequences of Han1 and CHM13 shows the high collinearity of every chromosome between the two genomes. (B) A gene order plot, where genes were numbered from 1 to N in order along all chromosomes of both CHM13 and Han1, and color-coded according to sequence identity. For both (A) and (B), the X axis is the CHM13 reference, and the Y axis is Han1. Chromosomes are in the order of 1 to 22, X, Y, and M. Alignments and dot plot produced using the MUMmer software, and gene order plot generated by Liftofftools.

**Figure 3.**
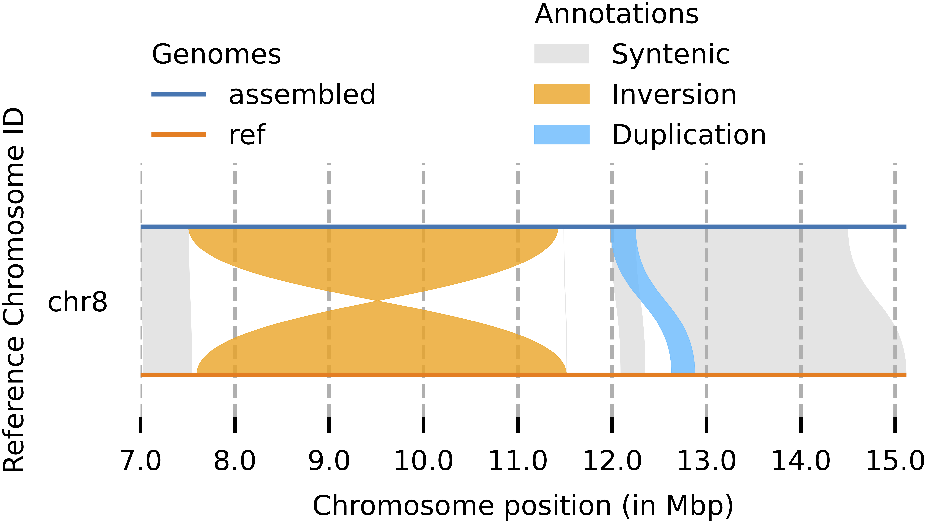
A detailed view of an inversion between Han1 (top, in blue color) and CHM13 (bottom, in orange color), spanning the region from approximately 7.5Mbp to 11.5Mbp on the chromosome 8. Figure created with Plotsr (Goel and Schneeberger 2022)

### Gene content differences between Han1 and CHM13

The recent completion of the genome of a single individual, CHM13, along with the results here for Han1, provided us with the opportunity to do the first-ever direct comparison between the gene content of two complete individual human genomes. In total, the Han1 annotation contains 60,708 genes including 20,003 protein-coding genes, 18,321 lncRNAs, and 658 rRNAs. The number of genes on Han1 is slightly lower than the 61,140 genes and 20,022 protein-coding genes on CHM13, although the small differences could be due to mapping artifacts. An overview of the comparison between Han1 and CHM13 by gene categories, along with a comparison to the gene content of GRCh38, is shown in Table 6.

**Table 6.** A comparison of major gene categories in the annotations of GRCh38, CHM13 and Han1

| Gene biotype | Number in GRCh38 | Number in CHM13 | Number in Han1 |
| --- | --- | --- | --- |
| protein coding | 19,871 | 20,022 | 20,003 |
| lncRNA | 17,793 | 18,389 | 18,321 |
| pseudogene | 15,357 | 16,027 | 15,881 |
| miRNA | 1,914 | 2,046 | 2,058 |
| transcribed pseudogene | 1,221 | 1,262 | 1,232 |
| rRNA | 38 | 765 | 658 |
| other | 2,506 | 2,629 | 2,556 |
| Total | 58,700 | 61,140 | 60,708 |

As described in Methods, we identified 46 distinct protein-coding genes that appear to have damaging mutations in both copies of the gene. Among these, 27 are frameshifts, 14 are 3’ truncations, 3 have lost their start codons, and 2 have gained an early stop codon. A list of the genes in each category appears in Supplementary Table S1. Out of these 46 genes, 24 are hypothetical proteins with no known function, raising the question of whether they are actually functional proteins. Six of these are olfactory receptors, which might indicate a genuine difference in the number of olfactory receptors, or might indicate that those copies are pseudogenes. Three are not actual genes but VDJ segments, which are highly diverse among individuals.

The remaining 13 genes are listed in Table 7. Two of these genes, MUC19 and AQP12A, contain premature stop codons and appeared to be severely truncated in Han1, with protein lengths far shorter than the corresponding proteins in both CHM13 and GRCh38.

**Table 7.**
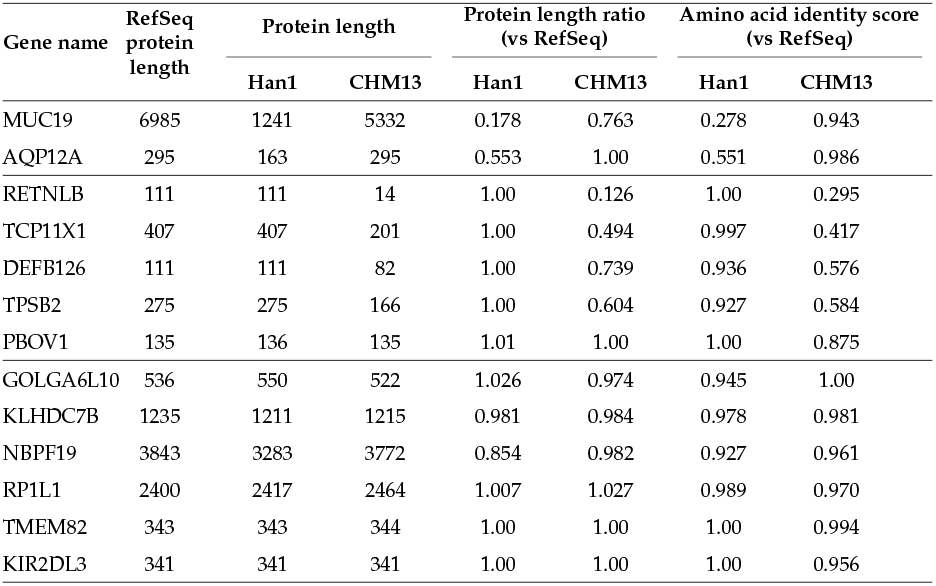
Protein level comparison between the Han1 and CHM13 genomes, showing genes with homozygous mutations in Han1 that are either frameshifted or truncated with respect to CHM13. The RefSeq protein length is based on the RefSeq v110 annotation of GRCh38.

| Gene name | RefSeq protein length | Protein length |  | Protein length ratio (vs RefSeq) |  | Amino acid identity score (vs RefSeq) |  |
| --- | --- | --- | --- | --- | --- | --- | --- |
|  |  | Han1 | CHM13 | Han1 | CHM13 | Han1 | CHM13 |
| MUC19 | 6985 | 1241 | 5332 | 0.178 | 0.763 | 0.278 | 0.943 |
| AQP12A | 295 | 163 | 295 | 0.553 | 1.00 | 0.551 | 0.986 |
| RETNLB | 111 | 111 | 14 | 1.00 | 0.126 | 1.00 | 0.295 |
| TCP11X1 | 407 | 407 | 201 | 1.00 | 0.494 | 0.997 | 0.417 |
| DEFB126 | 111 | 111 | 82 | 1.00 | 0.739 | 0.936 | 0.576 |
| TPSB2 | 275 | 275 | 166 | 1.00 | 0.604 | 0.927 | 0.584 |
| PBOV1 | 135 | 136 | 135 | 1.01 | 1.00 | 1.00 | 0.875 |
| GOLGA6L10 | 536 | 550 | 522 | 1.026 | 0.974 | 0.945 | 1.00 |
| KLHDC7B | 1235 | 1211 | 1215 | 0.981 | 0.984 | 0.978 | 0.981 |
| NBPF19 | 3843 | 3283 | 3772 | 0.854 | 0.982 | 0.927 | 0.961 |
| RP1L1 | 2400 | 2417 | 2464 | 1.007 | 1.027 | 0.989 | 0.970 |
| TMEM82 | 343 | 343 | 344 | 1.00 | 1.00 | 1.00 | 0.994 |
| KIR2DL3 | 341 | 341 | 341 | 1.00 | 1.00 | 1.00 | 0.956 |

Five genes, RETNLB, TCP11X1, DEFB126, TPSB2, and PBOV1, have protein sequences that are well-conserved compared to GRCh38, with 93–100% identity. These genes appeared to be frameshifted in CHM13, creating protein products in CHM13 that are either much shorter or frameshifted, with amino acid identity ranging from 30% to 88%. These results indicate that the damaging mutations occur in the CHM13 genome rather than in Han1.

For the other six genes, GOLGA6L10, KLHDC7B, NBPF19, RP1L1, TMEM82 and KIR2DL3, the translations of their CDS regions are well-conserved despite the mutations, with all of their amino acid identity scores higher than 92% and protein length ratios close to 1.

### Conserved synteny

Figure 2B shows the overall conserved gene order between the Han1 and CHM13 genomes. The figure shows 60,746 points, each representing a distinct gene, plotted in their order along the chromosomes. The color coding indicates the sequence identity score between CHM13 genes and Han1 genes, with green higher and red lower. Among the 60,746 genes, around 98.2% of genes (59,654) have sequence identity scores greater than 95%. As the figure shows, nearly all genes occur in exactly the same order between the two genomes.

### Gene family expansions

Liftofftools clustered all protein sequences using relatively strict criteria, which identified 19,653 clusters of near-identical proteins, of which 122 clusters had at least 3 genes in CHM13. 94 of the 19,653 clusters have copy number changes, with all changes being copy number losses in Han1.

We looked more closely at the TSPY gene family, which occurs as a large tandem array with interspersed repeats on chromosome Y in both CHM13 and Han1. Both genomes have 46 copies of TSPY, and CHM1 has 20 pseudogenes while Han1 has 24. Worth noting is that one copy of TSPY3 in CHM13, although annotated as a protein-coding gene, has a premature stop codon that creates a truncated 237 aa protein, whereas the same gene on Han1 is full length at 308 aa. Thus CHM13 appears to have lost one copy of the TSPY gene family.

In the 94 clusters with copy number loss, we found 46 clusters in CHM13 that had no corresponding genes in Han1, 43 clusters that lost a single gene copy, and 5 clusters that had lost more than one copy. The last groups (clusters that lost >1 gene copy) are shown in Supplementary Table S2.

## Conclusion

In this study, we assembled and annotated the first gap-free genome, Han1, from a southern Han Chinese individual. The genome has a total size of 3,099,707,698 bp and all 22 autosomes, the sex chromosomes X and Y, and the mitochondrial genome are assembled end-to-end with no gaps. The vast majority of the sequence is collinear with the CHM13 and GRCh38 genomes, although we did observe at least one novel mitochondrial insertion and a number of small-scale sequence duplications. Annotation of Han1 generated 60,708 genes, of which 20,003 are protein-coding genes. Finally, we conducted the first-ever direct comparison between the gene content of two complete individual human genomes, Han1 and CHM13 to determine how many genes are truly different between the individuals. This analysis revealed two protein-coding genes in Han1 and five genes in CHM13 that appear to be severely truncated in comparison to the full-length version of the protein.

## Data availability

All raw reads for HG00621 were generated and released by the Human Pangenome Reference Center, and are available as an AWS Open Data set from https://github.com/human-pangenomics/hpgp-data. The Han1 assembly (v1.0) is available from GenBank under accession JANJEX000000000, and on Github at https://github.com/JHUCCB/ChineseHanSouthGenome.

## Author contributions

KC, MP, and SLS designed the research. KC and AVZ performed assembly, scaffolding, and error correction. KC did the annotation. KC, AVZ, MP, and SLS analyzed the results. KC, AVZ, MP, and SLS wrote the manuscript. The authors thank Alaina Shumate for help with the rDNA annotation.

---

**Supplementary Table S 1.**
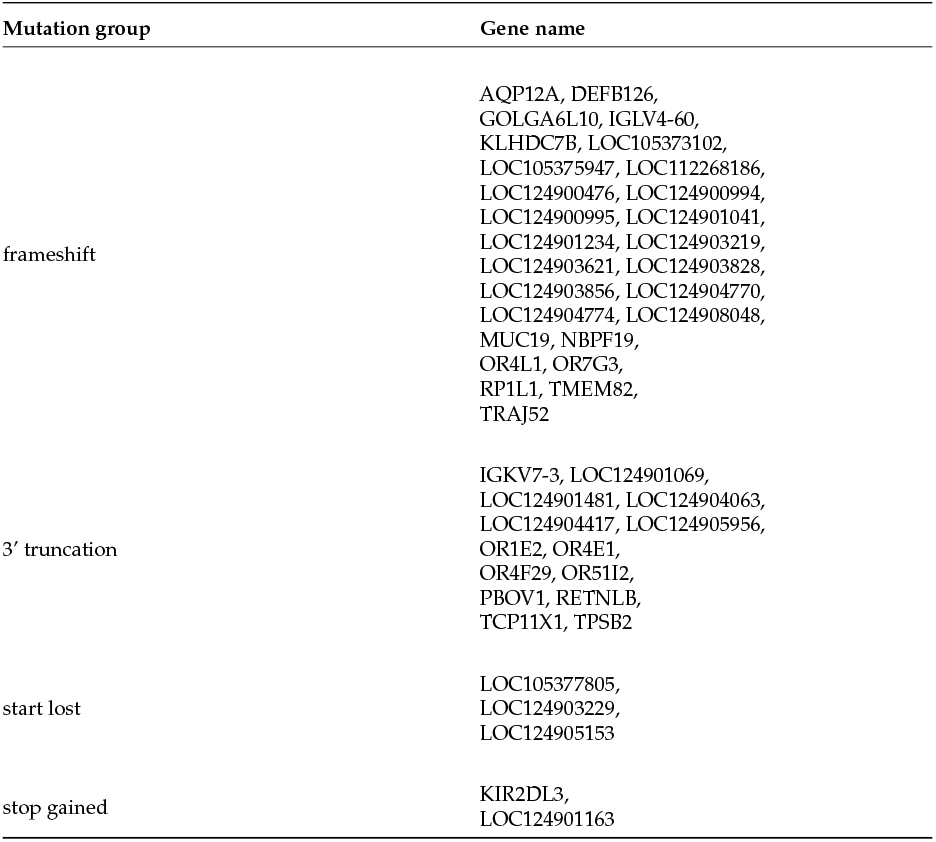
Genes that have at least 1 homozygous, non-SNP mutation in Han1 as compared to CHM13. Genes whose names begin with “LOC” are proteins with no known function. Genes whose names begin with “OR” are olfactory receptors. A frameshift is an insertion or deletion in the coding portion of the transcript that is not a multiple of 3 in length. A 3’ truncation denotes a truncated transcript where the lost sequence will produce a shorter protein. A “start lost” is either a truncation that deletes the 5’ end of the transcript including the start codon, or a mutation that changes the start codon to a different codon. A “stop gain” refers to a mutation that creates a stop codon, producing a shorter protein without necessarily altering the transcript length.

**Supplementary Table S 2.**
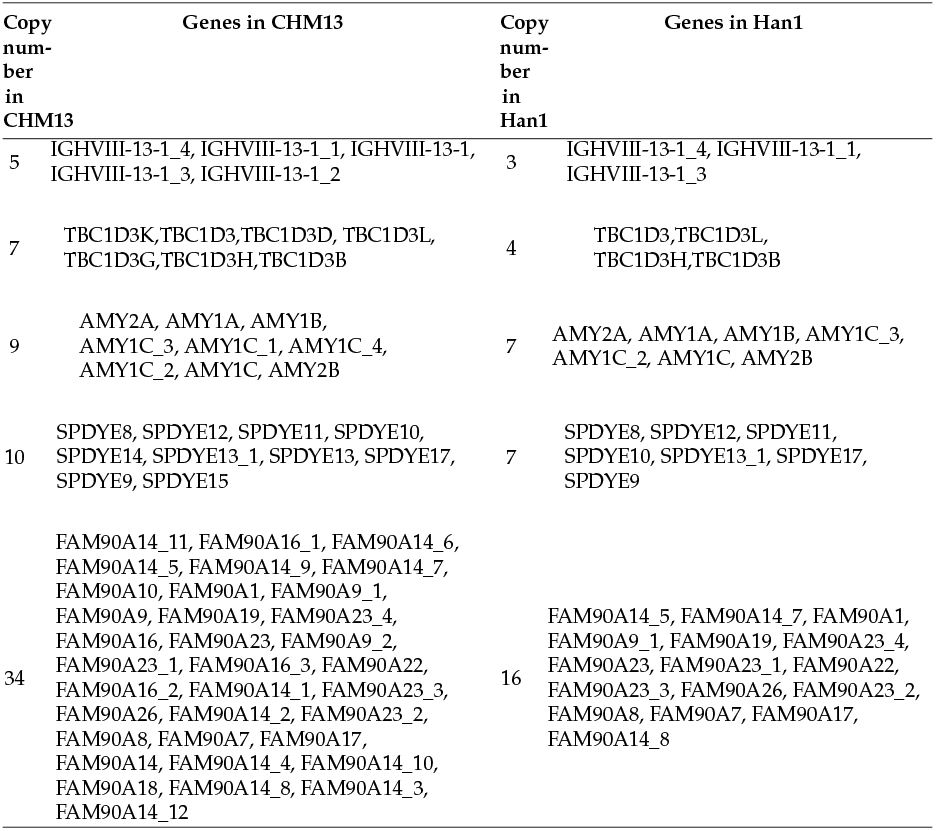
Genes that have more than 1 copy fewer in Han1 compared to CHM13.

**Supplementary Figure 1.**
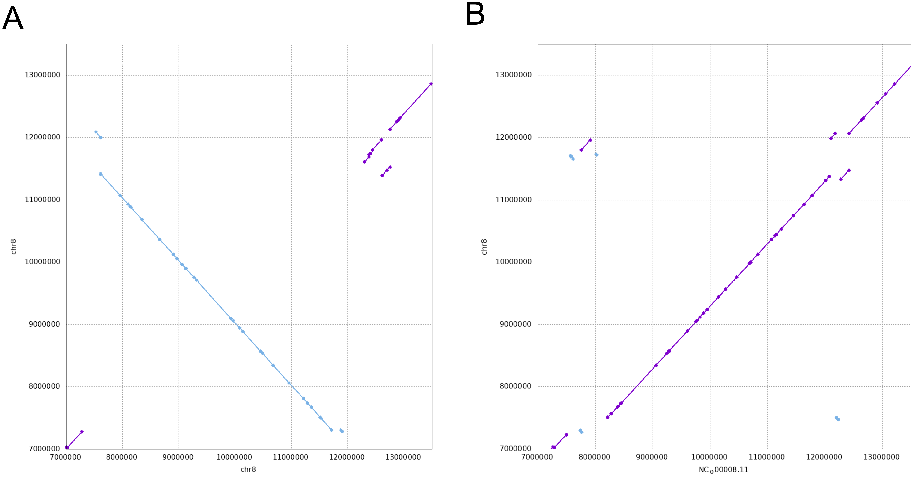
The zoomed-in dotplots on chromosome 8 from 7,000,000 to 13,500,000 showing the complex *β*-defensin gene cluster locus visualized by mummerplot. (A) demonstrates the inversion between CHM13 and Han1 in this region with CHM13 on the X axis and Han1 on the Y axis. (B) shows the collinearity between GRCh38 and Han1 in this region with CHM13 on the X axis and Han1 on the Y axis.

