## supplementary tables & figures for "The first gapless, reference-quality, fully annotated genome from a Southern Han Chinese individual"

**Supplementary Table S 1** Genes that have at least 1 homozygous, non-SNP mutation in Han1 as compared to CHM13. Genes whose names begin with "LOC" are proteins with no known function. Genes whose names begin with "OR" are olfactory receptors. A frameshift is an insertion or deletion in the coding portion of the transcript that is not a multiple of 3 in length. A 3′ truncation denotes a truncated transcript where the lost sequence will produce a shorter protein. A "start lost" is either a truncation that deletes the 5′ end of the transcript including the start codon, or a mutation that changes the start codon to a different codon. A "stop gain" refers to a mutation that creates a stop codon, producing a shorter protein without necessarily altering the transcript length.

| Mutation group | Gene name |
|---|---|
| frameshift | AQP12A,<br>DEFB126,<br>GOLGA6L10,<br>IGLV4-60,<br>KLHDC7B,<br>LOC105373102,<br>LOC105375947,<br>LOC112268186,<br>LOC124900476,<br>LOC124900994,<br>LOC124900995,<br>LOC124901041,<br>LOC124901234,<br>LOC124903219,<br>LOC124903621,<br>LOC124903828,<br>LOC124903856,<br>LOC124904770,<br>LOC124904774,<br>LOC124908048,<br>MUC19,<br>NBPF19,<br>OR4L1,<br>OR7G3,<br>RP1L1,<br>TMEM82,<br>TRAJ52 |
| 3′ truncation | IGKV7-3,<br>LOC124901069,<br>LOC124901481,<br>LOC124904063,<br>LOC124904417,<br>LOC124905956,<br>OR1E2,<br>OR4E1,<br>OR4F29,<br>OR51I2,<br>PBOV1,<br>RETNLB,<br>TCP11X1,<br>TPSB2 |
| start lost | LOC105377805,<br>LOC124903229,<br>LOC124905153 |
| stop gained | KIR2DL3,<br>LOC124901163 |

**Supplementary Table S 2** Genes that have more than 1 copy fewer in Han1 compared to CHM13.

| Copy number in CHM13 | Genes in CHM13 | Copy number in Han1 | Genes in Han1 |
|---|---|---|---|
| 5 | IGHVIII-13-1_4, IGHVIII-13-1_1, IGHVIII-13-1, IGHVIII-13-1_3, IGHVIII-13-1_2 | 3 | IGHVIII-13-1_4, IGHVIII-13-1_1, IGHVIII-13-1_3 |
| 7 | TBC1D3K,TBC1D3,TBC1D3D, TBC1D3L, TBC1D3G,TBC1D3H,TBC1D3B | 4 | TBC1D3,TBC1D3L, TBC1D3H,TBC1D3B |
| 9 | AMY2A, AMY1A, AMY1B, AMY1C_3, AMY1C_1, AMY1C_4, AMY1C_2, AMY1C, AMY2B | 7 | AMY2A, AMY1A, AMY1B, AMY1C_3, AMY1C_2, AMY1C, AMY2B |
| 10 | SPDYE8, SPDYE12, SPDYE11, SPDYE10, SPDYE14, SPDYE13_1, SPDYE13, SPDYE17, SPDYE9, SPDYE15 | 7 | SPDYE8, SPDYE12, SPDYE11, SPDYE10, SPDYE13_1, SPDYE17, SPDYE9 |
| 34 | FAM90A14_11, FAM90A16_1, FAM90A14_6, FAM90A14_5, FAM90A14_9, FAM90A14_7, FAM90A10, FAM90A1, FAM90A9_1, FAM90A9, FAM90A19, FAM90A23_4, FAM90A16, FAM90A23, FAM90A9_2, FAM90A23_1, FAM90A16_3, FAM90A22, FAM90A16_2, FAM90A14_1, FAM90A23_3, FAM90A26, FAM90A14_2, FAM90A23_2, FAM90A8, FAM90A7, FAM90A17, FAM90A14, FAM90A14_4, FAM90A14_10, FAM90A18, FAM90A14_8, FAM90A14_3, FAM90A14_12 | 16 | FAM90A14_5, FAM90A14_7, FAM90A1, FAM90A9_1, FAM90A19, FAM90A23_4, FAM90A23, FAM90A23_1, FAM90A22, FAM90A23_3, FAM90A26, FAM90A23_2, FAM90A8, FAM90A7, FAM90A17, FAM90A14_8 |

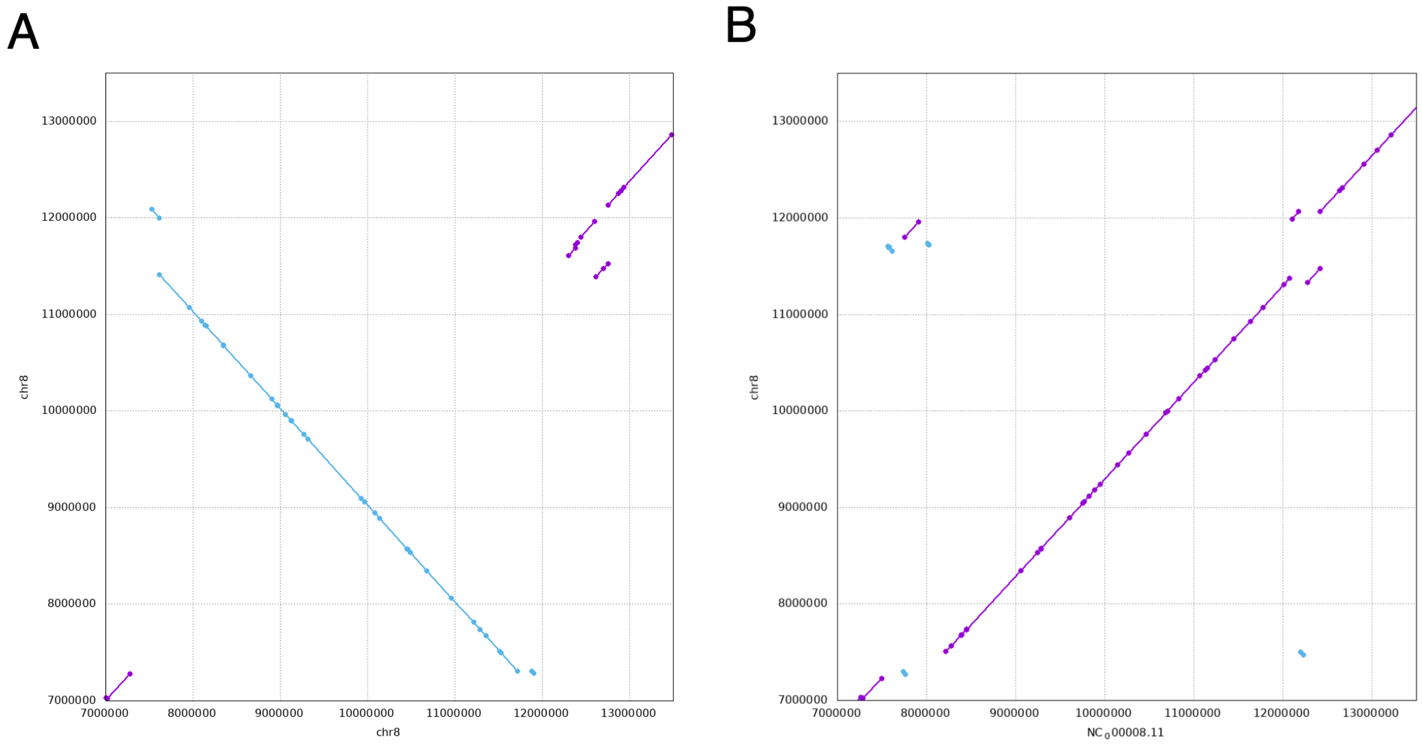


**Supplementary Figure 1** The zoomed-in dotplots on chromosome 8 from 7,000,000 to 13,500,000 showing the complex $\beta$-defensin gene cluster locus visualized by mummerplot. (A) demonstrates the inversion between CHM13 and Han1 in this region with CHM13 on the X axis and Han1 on the Y axis. (B) shows the collinearity between GRCh38 and Han1 in this region with CHM13 on the X axis and Han1 on the Y axis.
